# DNA metabarcoding of gut contents reveals key habitat and seasonal drivers of trophic networks involving generalist predators in agricultural landscapes

**DOI:** 10.1101/2022.05.07.489984

**Authors:** Hafiz Sohaib Ahmed Saqib, Linyang Sun, Gabor Pozsgai, Pingping Liang, Minsheng You, Geoff M. Gurr, Shijun You

**Affiliations:** State Key Laboratory for Ecological Pest Control of Fujian and Taiwan Crops, Institute of Applied Ecology, Fujian Agriculture and Forestry University, Fuzhou 350002, China; Guangdong Provincial Key Laboratory of Marine Biology, College of Science, Shantou University, Shantou 515063, China; Joint International Research Laboratory of Ecological Pest Control, Ministry of Education, Fuzhou 350002, China; Ministerial and Provincial Joint Innovation Centre for Safety Production of Cross-Strait Crops, Fujian Agriculture and Forestry University, Fuzhou 350002, China; BGI-Sanya, Sanya 572025, China; Azorean Biodiversity Group, Centre for Ecology, Evolution and Environmental Changes, University of Azores, Portugal; Southern Marine Science and Engineering Guangdong Laboratory, Zhuhai, Guangdong Province, 519000, China; Center for Infection and Immunity, the Fifth Affiliated Hospital, Sun Yat-sen University, Zhuhai, Guangdong Province, 519000, China; Graham Centre, Charles Sturt University, Orange, NSW 2800, Australia

**Keywords:** Lycosidae, high-throughput sequencing, food webs, trophic interactions, ecosystem services

## Abstract

**BACKGROUND:** Understanding the networks of trophic interactions into which generalist predators are embedded is key to assessing their ecological role of in trophic networks and the biological control services they provide. The advent of affordable DNA metabarcoding approaches greatly facilitates quantitative understanding of trophic networks and their response to environmental drivers. Here, we examine how key environmental gradients interact to shape predation by Lycosidae in highly dynamic vegetable growing systems in China.

**RESULTS:** For the sampled Lycosidae, crop identity, pesticide use, and seasons shape the abundance of preydetected in spider guts. For the taxonomic richness of prey, local- and landscape-scale factors gradients were more influential. Multivariate ordinations confirm that these crop-abundant spiders dynamically adjust their diet to reflect environmental constraints and seasonal availability to prey.

**CONCLUSION:** The plasticity in the diet composition is likely to account for the persistence of spiders in relatively ephemeral brassica crops. Our findings provide further insights into the optimization of habitat management for predator-based biological control practices.

## 1 INTRODUCTION

Understanding the predator-prey interaction networks and the analysis of processes shaping them are currently key objectives of ecology. These predator-prey interactions are critical in shaping the distribution, behavior, and abundance of prey and predator species within an ecosystem. Food webs can be used to comprehend both broad- and mesoscale distribution of species in these predator-prey interaction networks. Earlier predator-prey interaction networks were predicated on the assumption of a static structure, but recent ecological studies have highlighted the dynamic nature of trophic interactions across environmental and spatial gradients^1–3^. Thus, investigating the dynamics of predator-prey interactions across variable environments is key to understanding ecosystem functioning and, in turn, the effectiveness of biological control in agroecosystems ^4,5^.

Indeed, environmental variability, particularly in frequently changing agricultural environments, is likely to play a crucial role in determining the dynamics and the strength of predator-prey interactions^6,7^ but empirical knowledge is yet lacking^8–12^. Because these predator-prey interactions are the sum of often divergent responses of species to environmental filtering as well as of multi-level biotic interactions, predicting how environmental gradients drive these interactions is a great challenge for ecology and this is especially true in highly dynamic vegetable producing systems, where prey and predators are both small and have short life cycles.

Despite the complexities and difficulties associated with dietary studies, the information they provide is vital in revealing species interactions, which can be used to support efforts to efficiently manage and conserve on-farm biodiversity as well as to assess the capability of natural enemies of regulating important ecological processes (i.e. pest suppression)^13,14^. The environmental dependency of predator-prey interaction networks has already been reported in several studies showing the crucial impact of the local abiotic environmental variables on the structure and size of these networks^3,7,15^. However, for effective biological control, there is an urgent need for ecologists to unveil how landscape composition, differences in pesticide use and focal crop identity shape predator-prey interaction networks across different seasons in agroecosystems.

In recent years, DNA metabarcoding, which combines the amplification and sequencing of small sections of DNA from samples, has emerged as a promising and powerful method that, when paired with high-throughput sequencing (HTS), can reveal a wide range of trophic interactions in complicated food webs^14,16,17^. DNA metabarcoding together with HTS offers improvements in prey species detection in the gut and fecal samples over conventional DNA-based approaches or morphological identification of consumed taxa, which may fail to detect the full range of prey, for instance, soft-bodied species^18^. This approach is especially suitable for identifying the prey of invertebrate predators where morphological identification is usually impossible. Metabarcoding can resolve prey taxa to species level in systems where it would be hard to do so otherwise, such as in fluid-feeding invertebrates like spiders^19–21^. Thus, DNA barcoding paired with HTS approaches has changed our understanding of the dynamics in complex predator-prey interactions, eliminating many of the shortcomings of traditional methodologies.

Spiders are often abundant in agroecosystems and have a wide range of trophic interactions as biological control agents of pests and predators of non-pest prey ^21–23^. Though earlier studies have used molecular analysis of spider gut contents, there has been little attention to environmental drivers of prey range. Accordingly, we investigated how local factors (including pesticide use and crop identity) and the surrounding landscape composition, across different spatial scales and seasons, affect the assemblage structure of prey species of Lycosidae spiders in Brassica vegetable fields. Specifically, we hypothesized that (1) the prey assemblage structure is significantly affected by field-scale variables including crop identity and pesticide use, (2) responses of prey assemblages vary across seasons, and (3) increasing proportions of non-brassica land uses in the surrounding landscape influence the assemblages of prey species in the gut of spiders.

## 2 MATERIAL AND METHODS

### 2.1 Study system and data collection

We sampled spiders once in every season (i.e., spring, summer, autumn and winter) from Chinese cabbage and cauliflower farms located in Fujian Province, southeastern China in 2019 (Figure 1A) at the time of crop maturity. Smallholdings in this region are typically farms having highly dynamic vegetable growing systems, which are typical of those in other vegetable systems worldwide. A total of 18 fields at least 1 km apart were selected to represent different pesticide uses and crop identities with varying proportions of land use in the surrounding landscapes.

**Figure 1.**
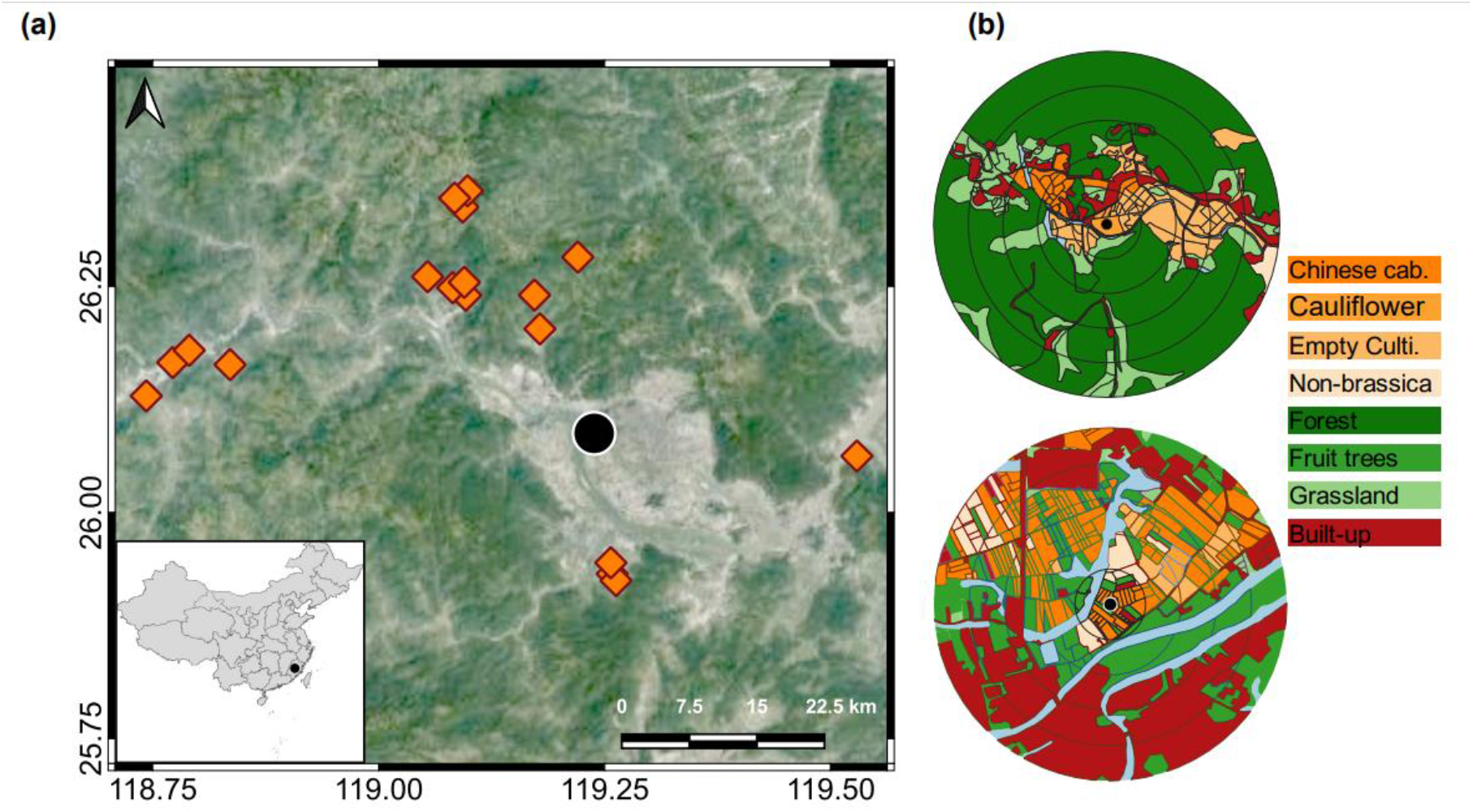
**(a)** Map showing the sampling locations in Fujian Province, southeastern China. **(b)** Example of landscape mapping of different habitats within 100, 200, 300, 400 and 500 m radius buffers around the focal sampling field.

Fields were divided into two groups based on management practices: 12 conventionally managed (i.e., synthetic pesticides or fertilizers were used) and six organically managed (i.e., no synthetic pesticides or fertilizers were used). Chinese cabbage and cauliflower were grown using direct sowing and seedling transplanting methods. We did not dictate the inputs on either conventional or organic farms nor did we intervene in any management practices. The disparity in field count between these two groups reflected their relative representation in this region. In all four seasons, a total of 93 Chinese cabbage (*Brassica rapa pekinensis*) and 151 cauliflower (*Brassica oleracea*) crop fields were selected which were well-distributed in both organic and conventionally managed fields.

To ensure that the leafy Brassica crops were not damaged and prevent surface DNA contamination, spiders were hand-collected using the sampling method 4 described in the studies of Sørensen et al. and Mader et al ^24,25^. Only adult spiders belonging to the Lycosidae family were hand-collected. Individual lycosids were sampled from the soil surface and plants directly into 5 mL clean tubes, by two people for an hour of searching per site. Samples were immediately transported to the laboratory in an icebox and then kept at −80 °C.

### 2.2 Landscape analysis

We used a drone (PHANTOM 4, Shenzhen Dajiang Baiwang Technology Co., Ltd., China) to take aerial images within a 500 m radius from each sample field. Images were used to classify the vegetation types into grassland, forest, built-up (e.g., residential land, greenhouses, and roads), water surfaces (e.g., small streams and ponds), Chinese cabbage, cauliflower, other Brassica crops (e.g., broccoli, canola, and mustard), non-brassica crops (e.g., pepper, eggplant, corn, and beans) and uncultivated (arable having no crop). QGIS 3.4 was used to calculate the proportions of various habitat types and field edge lengths (representing the total length of the field borders) in the 500-meter radius landscape surrounding the focal field, which was divided into five concentric buffer circles at intervals of 100-meters (Figure 1b).

### 2.3 DNA extraction, PCR and library preparation

A modified salt DNA extraction protocol (Sunnucks and Hales 1996^26^) was used to extract the genomic DNA of 732 adult spiders. Before performing DNA extractions, individual spiders were surface sterilized with ethanol and washed three times with double distilled water. Three individuals collected from the same fields were pooled to perform a single DNA extraction. All the genomic DNA was kept at −80 °C until it was needed.

To detect the barcode prey DNA, the PCR amplifications were performed using a previously developed primer pair designed for Lycosidae (NoSpi2)^27^, targeting fragments internal of the cytochrome c oxidase subunit I (COI) Folmer region^28^. To make the final PCR volume of 10 μl, each reaction contained 1 μl of template DNA (>50ng/μl), 5 μl KOD FX Neo Buffer, 2μl (2 mM) dNTPs, 0.3μl (10μM) each primer, 0.2 μl of KOD FX Neo DNA polymerase and 1.2 μl of ddH2O. The PCR was performed by initial denaturation at 95 °C for 300 seconds, followed by 25 cycles (denaturation for 30 seconds at 95 °C, annealing at 50 °C for 30 seconds, extension at 72 °C for 40 seconds), with a final extension of 300 seconds at 72 °C. To purify the successful PCR products, VAHTSTM DNA magnetic beads were used to remove primers, dimers, salts, and deoxynucleoside triphosphates (dNTPs). Equally molar DNA libraries were prepared for pair-end sequencing using an Illumina Novaseq 6000 (Illumina, Inc.) platform. Each batch of DNA extraction and PCR amplification was tested for possible contaminants in reagents with a negative control, both of which were negative.

### 2.4 Bioinformatics

Raw sequence data were primarily filtered by Trimmomatic (version 0.33)^29^ based on the quality of a single nucleotide. Identification and removal of primer sequences were performed by Cutadapt (version 1.9.1)^30^. Reads were demultiplexed, and paired-end reads were merged using USEARCH (version 10)^31^ followed by chimera removal using UCHIME (version 8.1)^32^. The high-quality reads generated from the above steps were used in further analysis. Cleaned sequences from all samples were then pooled and clustered into molecular operational taxonomic units (MOTUs) using USEARCH (version 10)^31^ over a 97% identity threshold. Taxonomic assignments for the remaining MOTUs were determined by blasting against the BOLD and NCBI databases. For species-level assignments, a match with 98% similarity was required in at least one of the two databases; 95% similarity was required for assignment to genus level, 90% for family level, 85% for order level and 80% for class level as a rough proxy^15^. Some ambiguous MOTUs that were generated by PCR, sequencing errors or blast errors were not used in the statistical analysis.

### 2.5 Data analysis

To compare the prey taxonomic composition in spider guts, all statistical analysis was conducted in R software (version 3.6.3)^33^. The vegan package was used to assess β-dissimilarities via NMDS (non-metric multidimensional scaling) ordinations based on Bray–Curtis dissimilarities using OTU–sample count data matrices^34^. In addition, a PERMANOVA (permutational multivariate analysis of variance), implemented as the adonis() function, was conducted to model the influence of local factors on the Bray-Curtis dissimilarity matrix using the vegan package after 999 permutations. Seasons and local factors (crop identity and pesticide use) were used as explanatory variables for the count dataset, which we used as a response variable. To determine if the explanatory variables had positive or negative relationships with each NMDS axis, the values of all variables were plotted against the NMDS scores.

A dbRDA model, based on Euclidean distances, was used to unveil the relationship between prey taxa abundances and richnessess in the spiders’ gut and environmental constraints (integrating both local and landscape). The community matrices were Hellinger transformed before performing the distance-based redundancy analysis (dbRDA). This transformation is often used in zero and one inflated community datasets because it downweighs variables with low counts and many zeros. To test the multicollinearity among all of the dbRDA model predictors, we used variation inflation factors (VIFs) test^35^. Because environmental factors with VIF > 10 had collinearity with other environmental variables, they did not significantly contribute to the model’s variance and were excluded from our final model. In the final dbRDA models of both abundance and richness only the spatial scale-related variables (i.e. radius size from the focal sample field) were found to be redundant (have VIF>10), so they were removed from the final dbRDA models. Additionally, an ANOVA-like permutation (999) test was used to determine the significance of RDA models^36^.

## 3 RESULTS

In total, 415 prey operational taxonomic units (OTUs) and 161333975 clean reads were identified from the sequencing data. The dominant prey orders were Diptera (123 species), Coleoptera (46 species), Hemiptera (28 species), Entomobryomorpha (20 species) and Lepidoptera (14 species). In terms of relative abundance, the diet of spiders mainly consisted of insects (79.72%). At the order level, 78% of the prey taxa detected in the gut of spiders were Coleoptera (33.70%), Diptera (19.40%), Lepidoptera (9.21%), Hemiptera (8.25%) and Entomobryomorpha (7.43%) (Figure 2).

**Figure 2.**
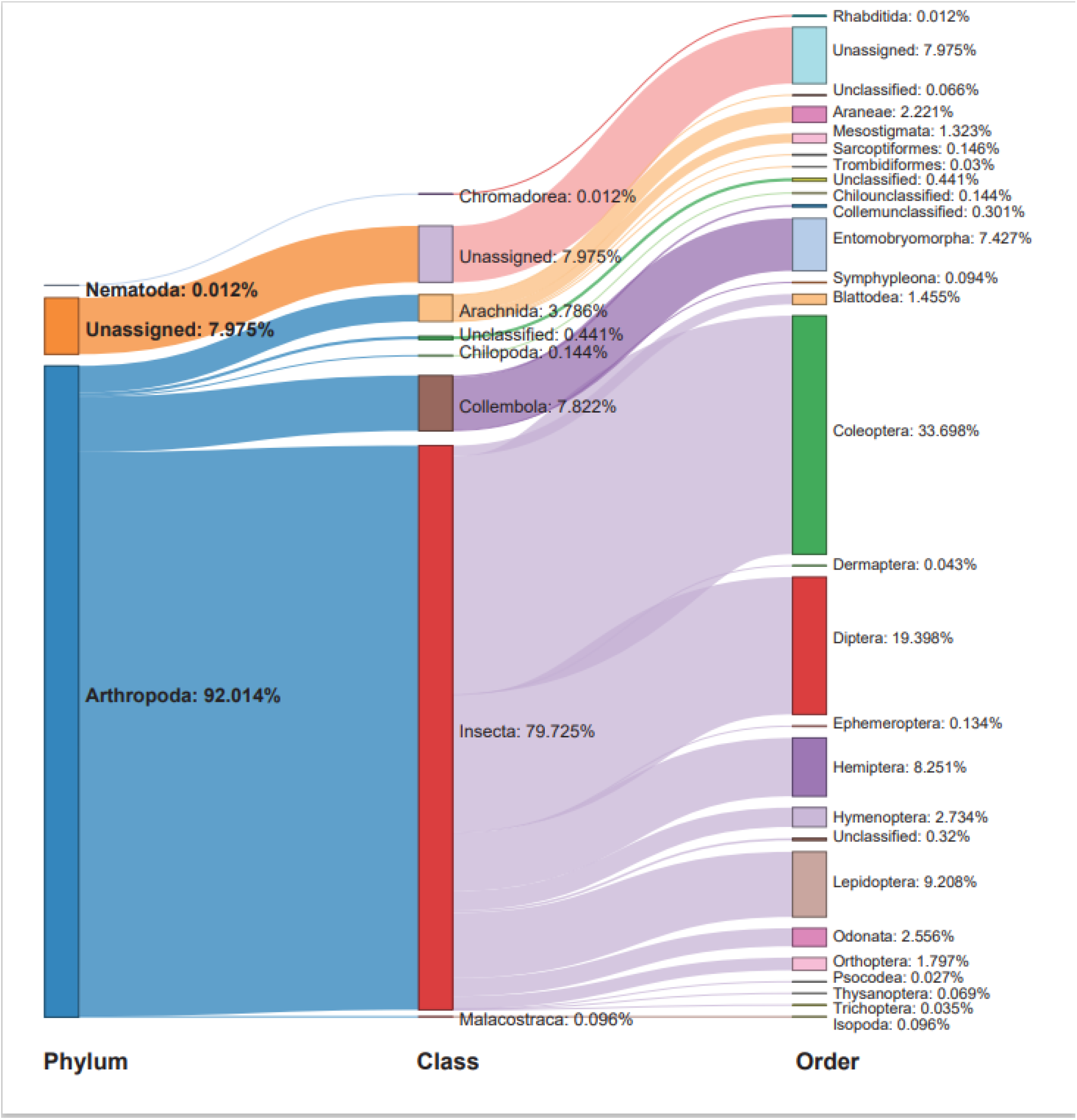
Sankey diagram of the relative abundances (%) of prey taxa detected in spiders’ guts.

### 3.1 Community structure across local factors and different seasons

Proportions of Coleoptera, Diptera, Hemiptera and Entomobryomorpha were relatively consistent across the four seasons, crop identity and pesticide use and mainly comprised of Bostrichidae, Drosophilidae, Anthocoridae and Entomobryidae, respectively. The predation of Lepidoptera by spiders was higher during the winter and autumn seasons, when gut content mainly comprised of Pyralidae and Plutellidae, compared with spring and summer seasons (Figure 3a–3d). Non-metric multidimensional scaling (NMDS) showed consistent differences in the community structure of prey species between crop identity and pesticide use across different seasons (Figure 4a–4b). Results of PERMANOVA were significant across both crop identity and pesticide use and their interaction with different seasons. The results of PERMANOVA supported the grouping in the NMDS ordinations (Figure 4a–4b).

**Figure 3.**
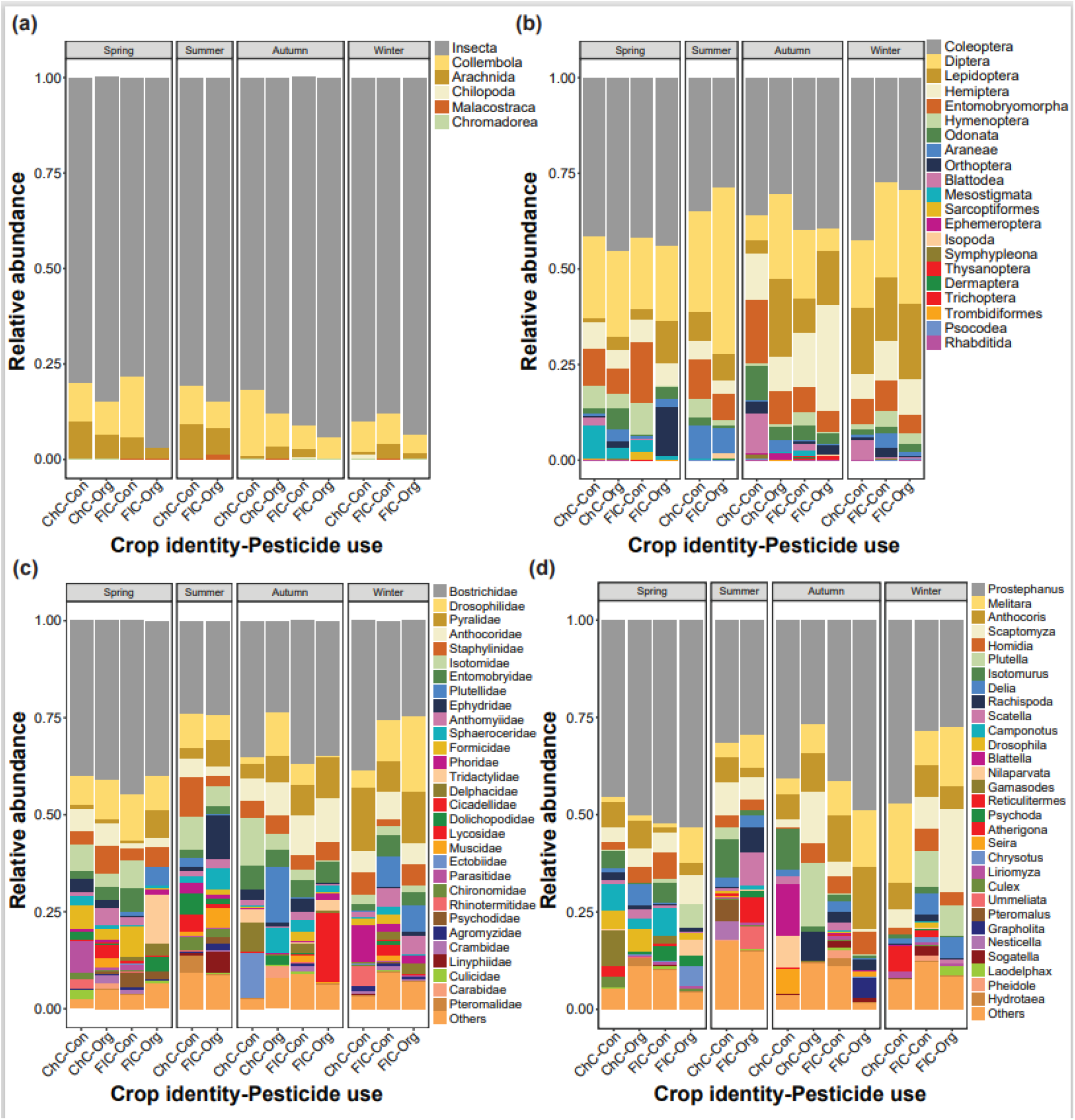
Relative abundance (%) of prey **(a)** classes, **(b)** orders, top 30 **(c)** families and **(d)** genera detected in the gut of spiders. Spiders were captured from different Brassica crop types (Chinese cabbage vs cauliflower) fields managed under different pesticide use (conventional vs organic) across four seasons. Here, “ChC”, “FlC”, “Con”, “Org”, “AUT”, “SPR”, “SUM” and “WIN” represent Chinese cabbage, cauliflower, conventional, organic, autumn, spring, summer and winter, respectively.

**Figure 4.**
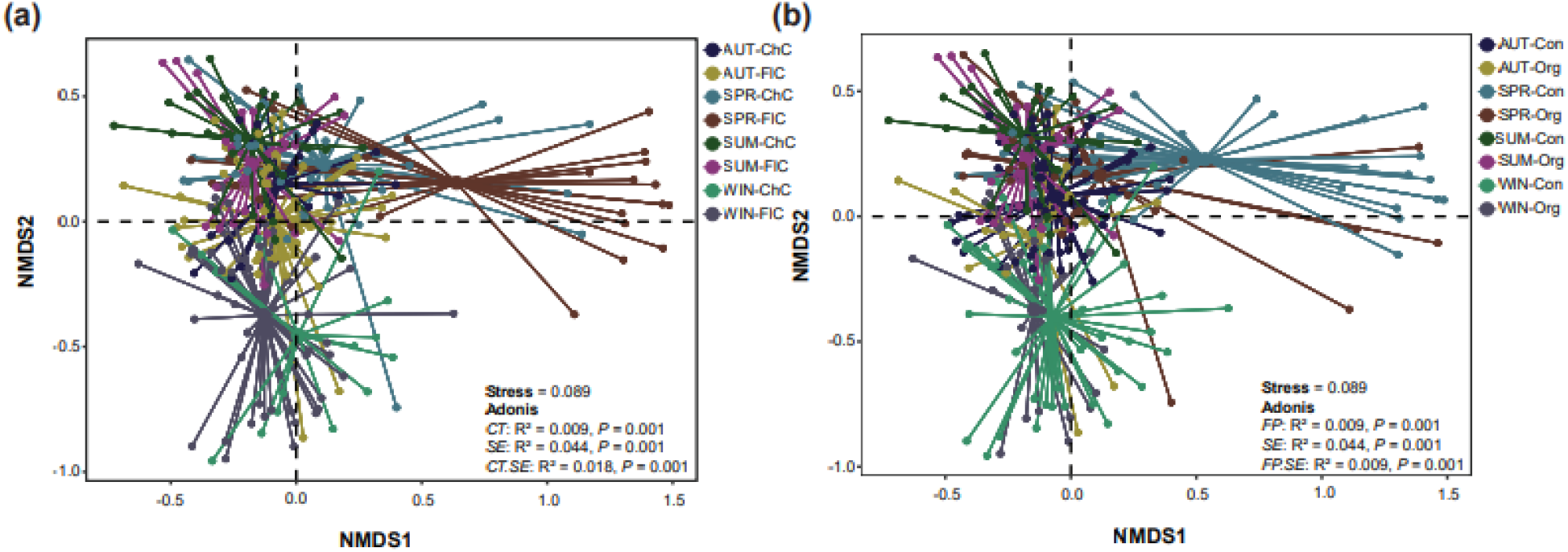
Plots of the non-metric dimensional scaling ordination based on the Bray-cutis dissimilarities for prey communities detected in the gut of spiders collected from **(a)** different Brassica crop identity (“ChC” = Chinese cabbage and “FlC” = cauliflower) and **(b)** different management systems (“Con” = conventional and “Org” = organic) across various seasons (“AUT” = autumn, “SPR” = spring, “SUM” = summer and “WIN” winter). Spider lines connecting each sample with the seasons-crop identity and season-pesticide use groups as centroid.

### 3.2 Local and landscape factors as determinants of prey species assemblage composition

The abundance and species richness of overall prey species and the top five orders showed inconsistent associations with landscape factors across different seasons, crop identity and pesticide use (Figure 5). The differences in the community structure based on the abundance of prey species detected in the gut of spiders were significant between local and landscape factors across different seasons (dbRDA model permutation test; *F* = 8.426, *p* = 0.001) and species richness (*F* = 9.529, *p* = 0.001). The first two axes in the dbRDA model explained 84% of the total variability in the assemblage structure of prey taxa based on their abundance and 65% based on their taxonomic richness in the diet of spiders. The edge density and elevation gradients only accounted for a small fraction of the variability in assemblages of prey species in terms of both abundance and richness. At the local scale, the assemblage structures of both abundance and species richness of prey taxa were significantly influenced by the Brassica crop identity, pesticide use, seasons, and elevation gradients (Table 1). The proportions of non-brassica crops and grassland patches in the surrounding landscape significantly explained the variability in the assemblage structure of both abundance and species richness of prey taxa (Table 1). The proportion of cauliflower had a significant influence on explaining the assemblage structure of prey, however only on species abundance-based data. On the other hand, proportions of Chinese cabbage, fallow land and water bodies had a significant contribution to explaining the assemblage structure of prey species in terms of their richness only (Table 1).

**Figure 5.**
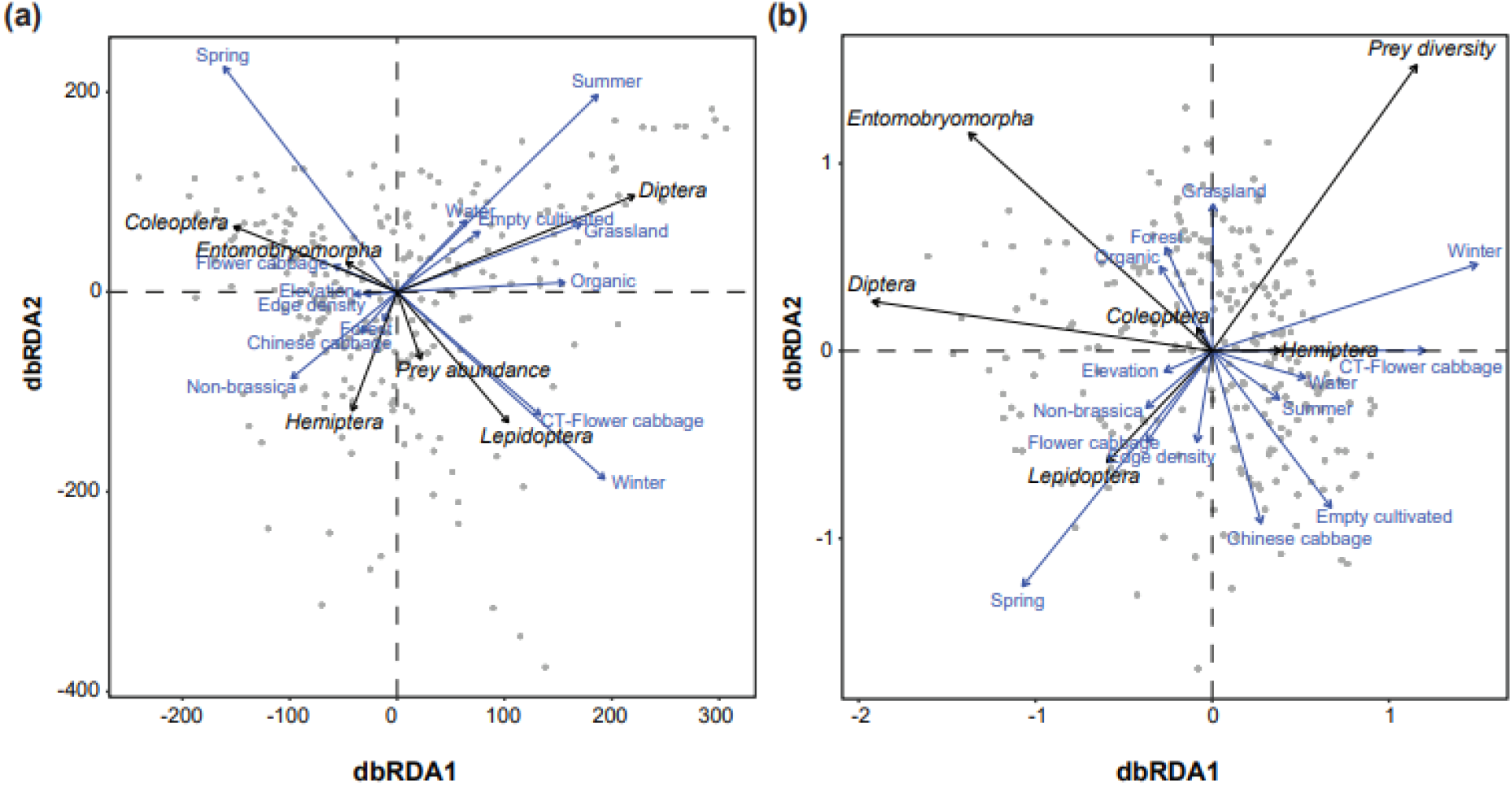
Distance-based redundancy analysis (dbRDA) illustrating the associations of prey taxa with various environmental factors in terms of **(a)** abundance and **(b)** richness in the diet of spiders. For each variable, the arrows’ length and orientation indicate the magnitude of explained variance. The association between prey taxa detected in spider guts and explanatory factors is represented by the perpendicular distance between them (below-90° = positive association and above-90°= negative association). The association is larger when the perpendicular distance is less.

**Table 1.**
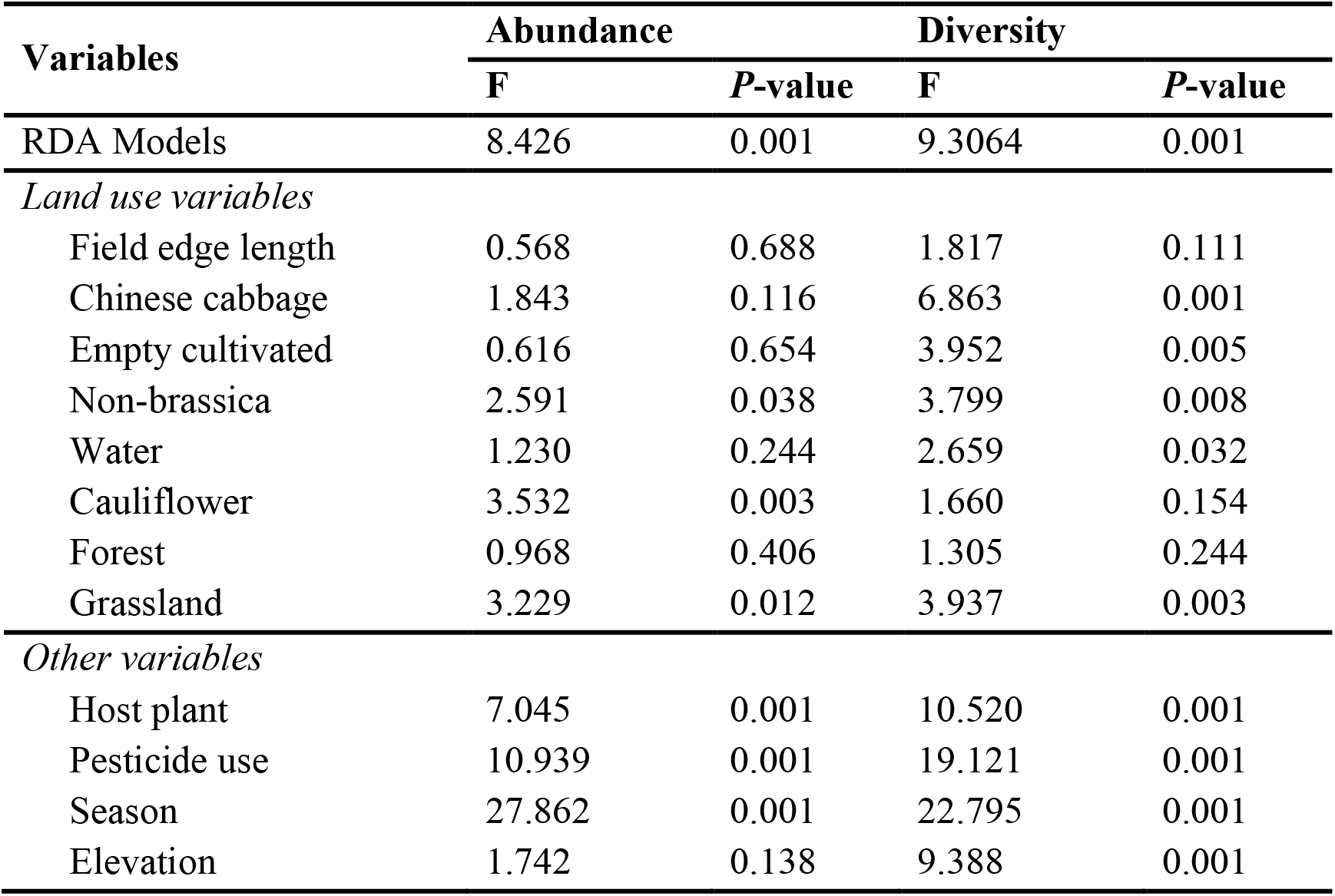
ANOVA table to test the significance of each predictor in the RDA model explaining the variance of prey communities in the gut of spiders.

The abundance of Diptera prey was higher during summer and the in organic fields with a high proportion of grassland, fallow land and water bodies. Conversely, the abundance of Coleoptera and Entomobryomorpha prey was strongly associated with a high proportion of cauliflower during the spring season. Cauliflower (as the focal crop identity) fields had a higher abundance of overall prey items and lepidopterans during winter. The higher abundance of hemipteran prey species was strongly linked to the fields with a higher proportion of forests, Chinese cabbage and non-Brassica crops in their surrounding (Figure 5a). For the species richness of prey taxa, cauliflower (as focal crop identity) and fields with a higher proportion of water bodies in the surrounding landscape had a greater diversity of overall prey species as well as hemipteran species during both summer and winter seasons. The organic fields with higher proportions of forest and grassland in the surrounding landscape had a higher richness of Entomobryomorpha, Diptera and Coleoptera prey species. Conversely, the assemblage structure of Lepidoptera prey species based on their species richness was strongly belonging to the fields with higher proportions of edge density, cauliflower, non-brassica crops and elevation gradients during the spring season (Figure 5b).

## 4 DISCUSSION

In this study, we identified how the trophic network structure of crop-dominant spiders dynamically adjusted with regard to the changing key environmental conditions and seasonality in a relatively ephemeral vegetable growing system. Furthermore, we concluded that the composition of spiders’ diet is eventually mediated, through the availability of prey species, by the gradients of local- and landscape-scale factors. We demonstrated how molecular data can be used for a novel purpose, to address changes in generalist predators’ interactions with prey and to investigate their drivers along different environmental gradients, particularly in ephemeral ecosystems where both predator and prey have a small size, a short life cycle and diverse foraging behaviors.

Beetles, flies, butterflies, moths, true bugs and collembola were the prey groups that accounted for 78% of all diet detected in the gut of Lycosidae. Overall, major groups of vegetable crop pests (including Chrysomelidae, Pyralidae, Plutellidae, Cicadellidae and Delphacidae) exhibited the larger fractions of spiders’ diet composition. The frequent detection of Diptera and Collembola along with small fractions of other prey taxa is likely explained by the spiders’ potential to utilize alternative, easily available, and highly nutritional prey (during the lower availability of major prey groups) to maximize their energy uptake to perform several metabolic activities, such as reproduction. Indeed, Diptera may have relatively high nutritional value and did not possess antipredator mechanisms except flight, which makes them optimal prey items. In addition, both Diptera and Coleoptera are highly active arthropods found in agricultural fields, which could also result in high encounter rates with predators, thus increased chances of predation by spiders. Similarly to our results, Binford et al., (2016) also reported >50% of *Tetragnatha eurychasma* Gillespie, 1992 (Aranea: Tetragnathidae) samples had Diptera prey in their jaws. Another study further described that Diptera occupy a very high proportion (>75%) of the total diet of *T. eurychasma*^38^.

Consistent with our expectations, the assemblage structure of arthropod prey in the guts of the spiders was influenced significantly by field-scale variables including crop identity and pesticide use across different seasons. The spiders’ diet in our study system was evidently enriched with additional arthropod prey species during spring and winter as compared to autumn and summer, regardless of the crop identity and pesticide use. In line with this, Roubinet et al., (2017) also showed substantial shifts in the diet of generalist predators (including spiders and carabids) between early and late cropping seasons. Indeed, in coastal ecosystems, the diet of spiders was shifted largely from a terrestrial diet in the spring and summer to the mostly marine diet in the late summer and autumn^40^. Variation among seasons is most likely explained by arthropod prey availability dynamics (i.e., variations in abundance and/or life stages of prey species), which are primarily determined by the available crops and cropping duration. Changes in arthropod prey community assemblages across different seasons most likely reflect the dietary adaptations to prevalent environmental factors, such as ambient temperature and humidity.

We observed that crop identity also acted as one of the key determinants of spider foraging behavior based on their dietary analysis. The guts of spiders captured from cauliflower crops had more abundant and diversified prey than Chinese cabbage. These patterns are also likely to reflect prey dynamics-driven changes in prey community assemblages, which are primarily determined by the presence or absence of the host crop^41^, its longevity^39,42^ and architectural characteristics^43–45^ and microenvironment quality^46,47^, and degree of disturbance^48–50^. A more diverse arthropod community may live on cauliflower crops due to its relatively longer cropping period and complex plant structure (which provides food and shelter). Additionally, the crops with long growing period can also offers a more stable microenvironment to support assemblages of diverse prey species. On the other hand, the shorter growth period, simpler plant structure, and higher frequency of harvesting and pesticide application events in the Chinese cabbage crops may explain the discrepancies in prey assemblages in the gut of spiders. A previous study has shown that simple habitat structure in young plants significantly decreases arthropod assemblages, compared to older plantations^39^. In this study, we found that highly disturbed growing systems with structurally simple crops have resulted in lower prey diversity in the spiders’ gut content, perhaps by limiting the resource availability for herbivores.

As the impacts of the host crops in this study, pesticide use patterns also had considerable effects on the prey assemblages in spiders’ diet; both overall abundances and richnesses of prey were significantly different in organically managed crops than in conventionally managed crops. It is widely accepted that increasing organic farming would lead to higher farmland biodiversity^51–53^. Plant species density was reported to be higher in organic fields (both inside the crop fields and adjacent non-crop areas)^54^ than in conventional ones, which may provide plenty of resources, and thus contribute to the more diverse assemblages of arthropod prey. Conventional farms utilize more synthetic herbicides^55^ to ensure weed control, resulting in lower plant species diversity (both within the crop fields and in adjacent areas), which ultimately has severe impacts on the diversity of inhabiting arthropod prey^22^. In addition, the availability of higher plant density and diversity outside the crop fields may provide alternative resources for several arthropod taxa to spill over into neighboring habitats^56–61^. Our findings further support that integration of organic schemes into vegetable growing systems can encourage more diverse assemblages of prey for predators. A diversified prey community can prevent the outbreak of a few dangerous pests, mostly because they can retain a high diversity of predators, which can promptly reduce and suppress the outbreak when a superabundance of a gradating pest occurs. Several studies have already suggested that higher biodiversity among natural enemies usually provides a better level of biological control^62,63^. In addition to this, the presence of non-pest prey, such as Entomobryomorpha, is likely to be important in terms of maintaining spider communities during times of low pest prey availability, but they will be killed by insecticides and indirectly suppressed by herbicidal weed control.

To date, gut content analysis studies have been only focused on relatively limited spatial scales, such as the detection of single or multiple prey species using conventional PCR methods^64^. Some recent dietary studies were focused on investigating the impact of a limited number of environmental variables (including elevation and vegetation mass) on a full dietary spectrum of generalist predators using DNA metabarcoding^15^. This study, however, shows the impact of different land-use variables along with previously described local factors on the full spectrum of predators’ diets at a landscape scale. Our results also showed that landscape composition variables can markedly affect the prey assemblages in the gut of spiders. The availability of semi-natural habitats (e.g. patches of grassland) and non-brassica crops (eg., pepper, eggplant, corn, and beans) in the surrounding landscape can significantly increase the abundance and prey species diversity^65^ in the gut of spiders captured on crop fields. These observed effects of semi-natural habitats and crops with longer duration were likely due to the availability of a stable understory habitat with a high diversity of plants that harbor alternative prey resources for generalist predators^66,67^. Indeed, several studies have also demonstrated that the availability of semi-natural habitat adjacent to crops with longer growing time can act as a strong factor in shaping arthropod assemblages^59,68–71^. Interestingly, the correlation between diet and landscape does not seem to be scale-dependent, we found no significant association between the landscape sampling radius and prey assemblages in spiders’ guts. A plausible explanation for this that is ground-dwelling Lycosidae rarely use ballooning and are likely to move frequently only between adjacent semi-natural habitats and the vegetable fields. Similarly, Schmidt et al., (2008) also reported that the lycosids exhibited no significant differences in their occurrences across varying spatial scales (ranging from 95-3000m), but the higher percentage of non-crop patches showed significant positive effects on lycosids. Therefore, the non-brassica crops and grassland patches available at shorter and longer distances from focal crop fields were the most influential for Lycosidae with limited dispersal ability but high mobility.

The ordination of prey taxa detected in the gut of spiders across different environmental variables highlighted distinct patterns of their association with surrounding land uses: in landscapes where the cauliflower was dominant, Lepidopta and Coleoptera prey were more common, whereas in non-brassica crops mostly Hemiptera (i.e., Delphacidae known as major pests of Asian rice systems) were consumed. A large amount of dipteran prey was associated with landscapes dominated by semi-natural habitats. These distinct patterns of prey association with different land use in the surrounding landscape can likely be explained by the spill-over (from focal crop to adjacent non-crop vegetations) and active hunting behavior of Lycosidae.

## 5 CONCLUSION

In conclusion, we demonstrate strong dietary differences of prey in the gut of spiders across key environmental variables. We used molecular metabarcoding approaches, targeting the short fragment of COI, which can offer the detection of a broad range of prey in the spiders’ gut. This method could be used to rapidly evaluate anthropogenic effects on biodiversity and ecosystem functioning, particularly in extremely dynamic environments such as vegetable plantations or other annual crops. We showed that crop-abundant spiders adapt their foraging behavior dynamically in response to environmental constraints and the seasonal availability of prey. The plasticity in the diet composition indicated that spiders are extremely adaptable to changing environmental conditions, implying their significance as biological control agents in agro-ecosystems, particularly those with highly dynamic growing systems. These findings indicate the conditions under which the biological control exerted by this group of spiders is most likely to be optimal and which habitat management efforts should strive to achieve.

## ACKNOWLEDGMENTS

This work was financially supported by the National Natural Science Foundation of China (No. 31972271), State Key Laboratory of Ecological Pest Control for Fujian and Taiwan Crops, Joint International Research Laboratory of Ecological Pest Control, Fujian-Taiwan Joint Innovation Centre for Ecological Control of Crop Pests, International science and technology cooperation and exchange program of FAFU (KXb16014A), and the Thousand Talents Program and the “111” Program in China.

## AUTHOR CONTRIBUTIONS

H.S.A.S., M.Y., S.Y. and G.M.G. conceived and designed the experiments. H.S.A.S. and L.S. conducted field sampling and lab experiments. H.S.A.S. performed the data analysis, prepared figures and tables. H.S.A.S., G.P. and G.M.G. interpreted the data and wrote the paper. P.L., S.Y. and G.P. assisted in the data analysis. All authors revised the final version and gave their approval for submission.

## DATA AVAILABILITY

All data supporting the results of this article are included in the published article and its supplementary information files.

